# ONTOLOGY™: Streamlined DNA Barcoding and Metabarcoding on Nanopore

**DOI:** 10.64898/2026.11.23.768700

**Authors:** Paul DN Hebert, Ken A Thompson, Robin M Floyd, Saeideh Jafarpour, Sean WJ Prosser, Suz Bateson, Nicholas W Bard, Emine Ozsahin

## Abstract

Nanopore sequencers have unlocked the potential for a distributed network of DNA barcoding and metabarcoding laboratories. However, the realization of this potential confronts two challenges—one molecular, one computational. First, rigorous indexing protocols are critical to correctly associate sequences with their source specimen or sample; and second, data interpretation often demands processing millions of indexed reads in a structured fashion via complex bioinformatic pipelines. ONTOLOGY™ addresses these challenges. At its core lies a graphical user interface (GUI) that guides users through the workflow from specimen or sample reception to the generation of DNA barcode records or metabarcode files. To facilitate data acquisition, it includes freeze-dried PCR plates that uniquely tag the amplicons from each well, hence supporting sample-sequence association. The ONTOLOGY GUI is constructed around two core workflows that process data from a sequencing Run. The <u>Barcode Inference Pipeline</u> (**BIP**) delineates the OTU(s) recovered from each well and designates them as target or non-target sequences via comparison of OTU sequences to a reference sequence library. The <u>Metabarcoding Analysis Pipeline</u> (**MAP**) processes sequence arrays derived from bulk samples (organismal or environmental DNA) and infers the species represented while rigorously excluding false positives. To validate ONTOLOGY, we present an original data analysis completed via both pipelines. Built for maximum flexibility, ONTOLOGY also supports the analysis of custom experiments. By employing a GUI to coordinate passage through the workflows with structured amplicon tagging and informatics pipelines, ONTOLOGY facilitates the molecular analyses needed to advance biodiversity science via decentralization. Plates and software can be obtained from www.ontology.bio.

## BACKGROUND

Over the past 20 years, DNA-based identification systems have gained broad adoption for basic and applied science, reflecting their effectiveness in specimen identification and species discovery [1–4]. For most of this interval, sequencing has been undertaken in core facilities because it required expensive instrumentation, but low-cost nanopore sequencers have shifted the situation [5, 6]. Although it is no longer alone [7], Oxford Nanopore Technologies (ONT) is the market leader. Its sequencers and their companion flow cells are well suited to support both DNA barcoding and metabarcoding. Its MinION sequencer uses two flow cells, the Flongle (to be discontinued February 2027) and MinION, which generate about 500,000 and 10 million 750 bp reads, respectively, while its P2 series sequencers support the use of PromethION flow cells that generate >100 million reads. As such, the MinION supports highly multiplexed barcode workflows [8] and some metabarcoding applications [9], while the PromethION delivers the sequencing depth required for metabarcoding [10].

Although nanopore sequencers are widely available, applying them for biodiversity research confronts two challenges: (i) the need for molecular indexing (i.e., tagging) systems to discriminate the sequences derived from each specimen/sample, and (ii) specialized informatics pipelines to analyze the resultant data. The need for an indexing system arises because amplicon sequencing on nanopore is expensive unless many specimens or samples are pooled. For example, sequencing 100K specimens on a MinION flow cell reduces the per-specimen sequencing cost to $0.01 [8]. While the design and deployment of unique molecular index (UMI) schemas to support highly multiplexed workflows is routine for core facilities [11], it represents a challenge in a distributed context because it requires technical expertise and considerable funding, and gains efficiency with specialized dispensing equipment. Once indexed sequences have been obtained, a bioinformatic pipeline is essential to connect sequences to their source specimen and to suppress off-target sequences. A well-structured pipeline is key for analyzing the 0.5–200M reads in a run [12], but with few exceptions [13–15] pipelines are tailored to researchers with bioinformatics training who are comfortable with a command-line interface [16].

ONTOLOGY™ is a new system for DNA barcoding and metabarcoding that trivializes the most complex parts of the workflows by integrating an amplicon indexing system with software that interprets the resultant sequence data. It is built around ONTOLOGY PCR kits and ONTOLOGY software. The PCR kits include 96-well plates provisioned with a standard PCR cocktail augmented by the UMIs needed to differentially tag the amplicons from each well in a single (i.e., ‘one-step’) PCR cycling reaction. As such, these plates solve the tagging challenge, allowing sequencing to proceed with varied levels of multiplexing. The plates are freeze-dried following their manufacture, enabling their global distribution without a cold chain. ONTOLOGY software employs an intuitive graphical user interface (GUI) to enable novices to execute two workflows to support data analysis using a reference sequence library needed for data validation and interpretation (e.g., taxonomic assignments). The DNA barcoding workflow uses BIP [17] to assemble OTUs and discriminate the barcode sequence from non-target sequences. The metabarcoding workflow uses MAP [10] to determine the source species for the sequences recovered from each bulk sample. ONTOLOGY workflows require only basic molecular biology skills—trainable in a short course—and no coding or use of the command line.

As ONTOLOGY PCR kits currently support DNA barcoding and metabarcoding using the standard COI barcode region for the animal kingdom via the AR-01 kits, this paper discusses its capabilities when analyzing this gene target. However, it can support the use of any gene region employing any primer set, and ONTOLOGY kits will soon be available for the barcode regions employed for other major taxonomic groups. This paper begins by describing the overall functionality of ONTOLOGY before introducing its solutions for indexing and data analysis. It then considers results gathered by Demonstration Runs before discussing implications for the establishment of a distributed network of analytical facilities.

## IMPLEMENTATION

There are two main components to ONTOLOGY: (i) freeze-dried PCR plates to generate tagged amplicons (with an optional DNA extraction add-on), and (ii) a downloadable GUI to orchestrate passage through the DNA barcoding (BIP) and metabarcoding (MAP) workflows. This section begins by describing the ONTOLOGY PCR plates and indexing system, then considers the role of its GUI in structuring passage through both workflows. It finishes by describing a joint barcode/metabarcode study that demonstrates ONTOLOGY’s capabilities. As BIP and MAP are already on preprint servers [10, 17], they are described only briefly here; interested readers can access full details in the preprints.

### 1. ONTOLOGY PCR Plates

Each ONTOLOGY kit includes a batch of freeze-dried [18] plates [19, 20]. The PCR cocktails in these plates are used to generate tagged amplicons and are stable at room temperature for two months and at 4 °C for at least a year (**Figure S1; Supplementary Information**). As such, their use requires only the addition of molecular grade water and template DNA. PCR kits can be used with both ONT and Qitan sequencers [7] without any modification to standard library preparation protocols.

***DNA Barcoding***: Each plate supports the analysis of 95 specimens plus one negative control, and all plates within a batch can be pooled in the same sequencing Run. Standard protocols involve the inclusion of one negative control in well H12. The sequencing cost to secure a specimen’s barcode declines as the number of plates used to create the amplicon pool rises. Regardless of the number of plates used, adequate sequencing depth can be achieved with a Flongle (ONT; **Figure S2**). A single QCell-384 (Qitan) or MinION (ONT) flow cell can support multiple Runs if washed and properly stored [20].

***DNA Metabarcoding***: Each plate can analyze up to 96 bulk samples, but standard protocols call for technical replication plus controls. Presuming three replicates per sample, a plate can support the analysis of 30 samples plus six controls (positive and/or negative). ONTOLOGY’s standard protocols suggest using one plate per PromethION flow cell to reach a target of ≥1M reads per technical replicate (**Figure S2)**. Since plates are not pooled in this design, ONTOLOGY’s symmetrical PCR kit can be used to improve read retention above that achieved with the asymmetrical AR-01 kits.

***Plate Identifiers***: Each batch of plates possesses a unique six-character alphanumeric code (e.g., AAA001) while the individual plates in it are discriminated by a two-digit suffix. For example, the sixth plate in batch AAA001 is AAA001-06.

***Primers***: At launch, ONTOLOGY kits are available only for COI via the AR-01 primer set, which targets arthropods. Although AR-01 achieves the highest recovery for arthropods, it is also very effective for many other animal lineages (Supplementary Information).

***Primer Tags***: Each primer in the AR-01 set is tagged with a 16 bp UMI with a different pair in each well. Each kit includes 10 plates whose 960 wells generate uniquely tagged amplicons by using 10 reverse primers and 96 forward primers [21]. The same reverse primer is shared by all wells in a plate but is unique to that plate. By contrast, a different forward primer is used for each of the 96 wells in a plate, but the same array is shared by all plates (e.g., well A01 in plate 1 has the same forward primer as A01 in plate 10). In the symmetrical plates, the same UMI is used as forward and reverse, allowing demultiplexing when only one is read. Further details on UMIs are provided in the <u>Supplementary Information</u>.

### 2. ONTOLOGY GUI

ONTOLOGY software provides a GUI that guides users through the three stages required to generate barcode or metabarcode data: 1) Prepare samples or specimens for analysis; 2) Implement DNA extraction, PCR, and sequencing; and 3) Analyze sequence data. Upon opening ONTOLOGY, the user selects a workflow: either <u>DNA Barcoding</u> (single specimens, **Figure 1**) or <u>Metabarcoding</u> (bulk samples).

**Figure 1:**
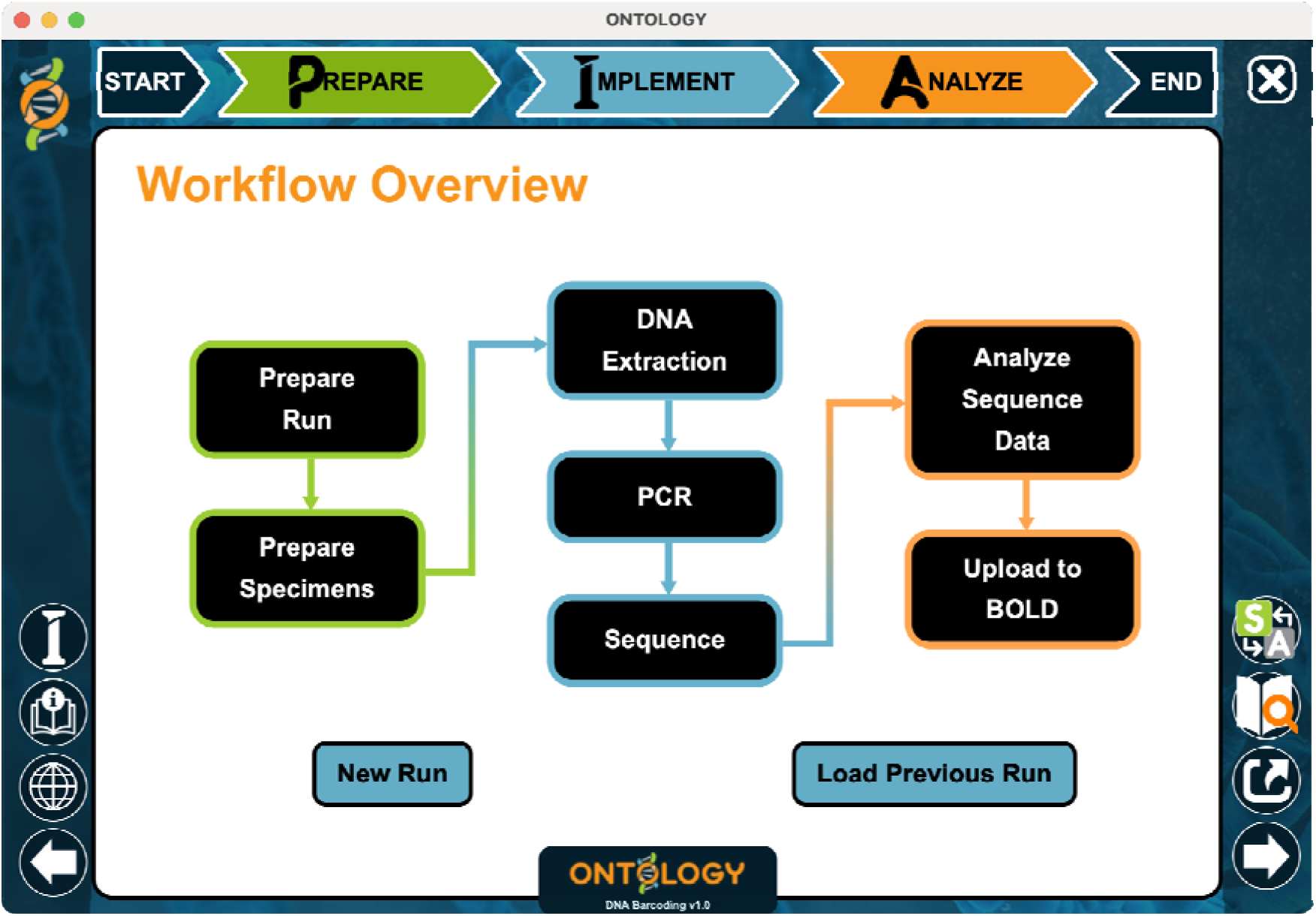
Overview of the key steps in ONTOLOGY’s DNA barcode workflow (Standard Mode).

Both workflows are organized around a Run: the set of specimens or samples that are destined for analysis on a single use of a flow cell. The number of PCR plates used in a Run is flexible, but the barcode workflow is most cost-effective when amplicons from all ten PCR plates are pooled. By contrast, metabarcoding studies typically employ a single plate because considerable read depth is required to maximize species recovery. Because sequencers and flow cells have differing error profiles, it is critical to record the technology employed in data acquisition [22]. ONTOLOGY meets this need by allowing the user to submit standard files that record the sequencer, flow cell, and base caller employed in each Run.

#### DNA Barcoding

The DNA barcode workflow (**Figure 1**) can be completed in Standard or Advanced mode. Standard mode guides a novice user through the entire workflow, while Advanced mode provides access to a simplified workflow initiated after a user has completed sequencing. Both modes lead to the recovery of the target DNA barcode for each specimen that contributed a DNA extract for analysis. The amplicons from each extract gain a unique UMI pair during PCR, and the library created by pooling samples can include amplicons from any number of plates in a batch.

In Standard mode, the DNA barcode workflow is as follows:

<u>Stage 1 – Prepare</u>: The workflow begins with the user entering the Batch ID and confirming the primer set (e.g., AR-01). The user is then guided through the six steps which require completion to initiate a barcode Run. These include: 1) Sort specimens; 2) Array them in a set of 95 plus one negative control; 3) Enter metadata; 4) Photograph; 5) Tissue sample; and 6) Label specimens. After these steps are completed for the first plate, they can be repeated for additional sets of 95 specimens by looping through Prepare. An interactive summary page details the specimens carried into the Run.

<u>Stage 2 – Implement</u>: This segment of the workflow guides progress through the three wet lab steps: 1) DNA extraction; 2) PCR; and 3) Library preparation and sequencing. The GUI provides detailed protocols for each step while the kit includes the consumables required to prepare DNA extracts as well as the PCR plates which generate indexed amplicons (**Figure 2**). Interactive calculators facilitate library preparation.

**Figure 2:**
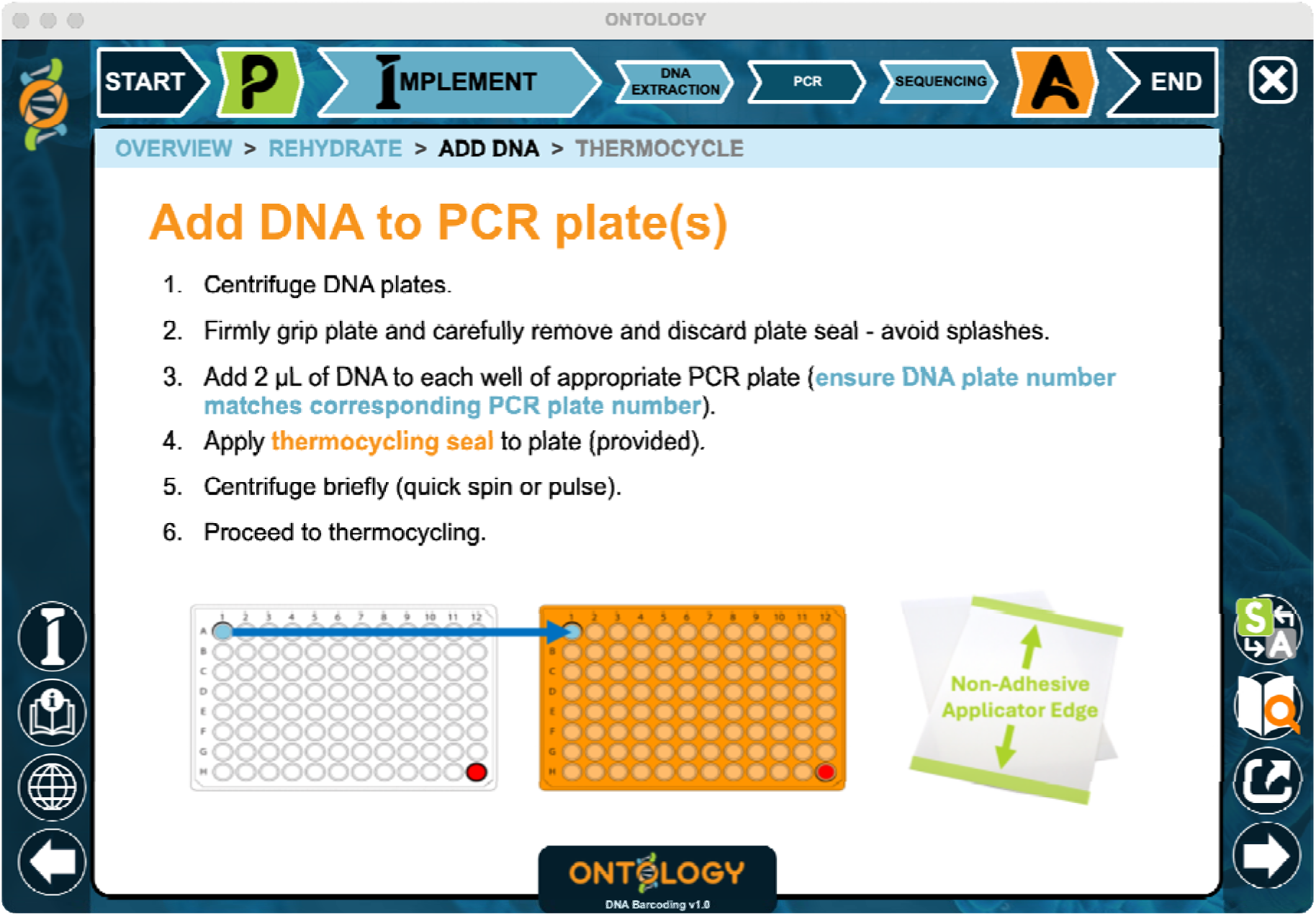
Screenshot from the Implement Stage of ONTOLOGY’s Barcode workflow (Standard Mode).

<u>Stage 3 – Analyze</u>: The Barcode Inference Pipeline (BIP) guides progress through the five steps required to analyze the sequence data: 1) Import sequence data; 2) Import or select reference library; 3) Perform bioinformatic analyses (**Figure 3**); 4) Evaluate technical/biological summaries; and 5) Upload data to BOLD [23, 24]. Prosser et al. [24] and the <u>Supplementary Information</u> provide further details on BIP. Bioinformatic parameters can be adjusted by the user if they wish to depart from defaults.

**Figure 3:**
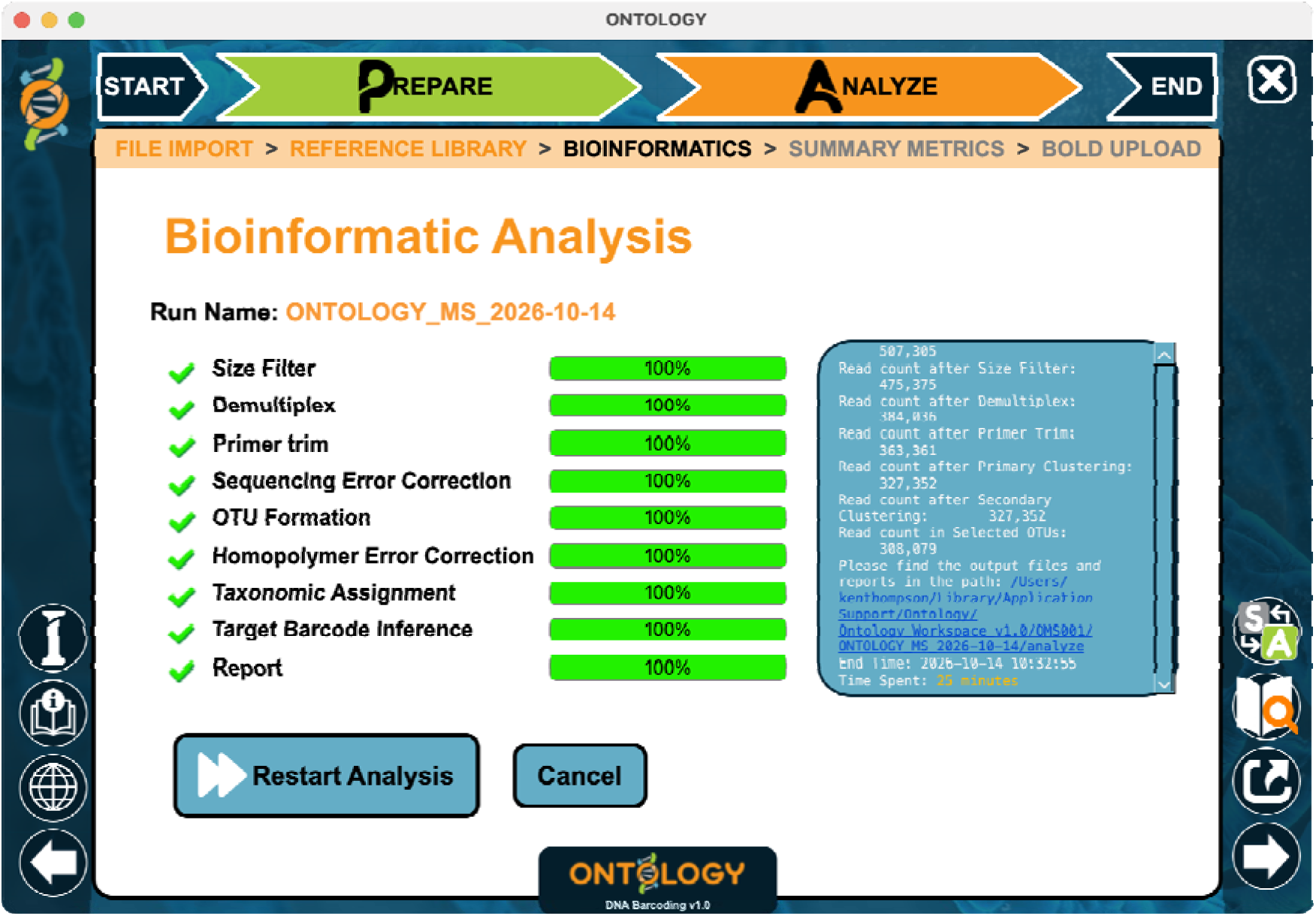
The analytical cascade in the Barcode Inference Pipeline (BIP) which transform the input.fastq.gz file(s) into a table of OTUs for the samples before generating a summary of the success in barcode recovery. The 10-plate Run was completed in Advanced mode using an M4 Pro MacBook Pro with 24 GB RAM running macOS Sequoia 15.6.1.

In Advanced mode, the Prepare Stage is greatly simplified with all information entered on a single page; the summary page is unchanged. Implement is not included (but can be accessed via a pop-up window) since sequencing has already been completed. Analyze is unchanged.

#### Metabarcoding

Structured similarly to the Advanced mode of the DNA barcoding workflow, the metabarcoding workflow is designed to ascertain the species present in bulk samples of air, soil, water, or organisms. DNA extracted from each sample is amplified and indexed during PCR and the amplicons from wells are pooled to construct a library. Species recovery depends on many factors, but read depth is key. The metabarcoding workflow includes the same Stages as DNA Barcoding, but protocols differ.

<u>Stage 1 – Prepare</u>: The metabarcoding workflow begins with the user entering the Batch ID and confirming the marker (COI) and primers. Like DNA barcoding’s Advanced mode, Prepare only requires entering metadata. Because samples destined for metabarcode analysis typically include multiple technical replicates plus positive and negative controls, the metadata sheets are specialized to accept this information. This well-to-well treatment variation in each metabarcoding plate needs to be registered during Prepare to guide subsequent data analysis.

<u>Stage 2 – Implement</u>: ONTOLOGY does not include reagents for the preparation of DNA extracts for metabarcode analysis because the optimal protocol depends upon the type of sample being analyzed (e.g., the protocol for bulk samples of organisms differs from that used to extract DNA from water). Following PCR, which typically involves fewer cycles for metabarcoding than for DNA barcoding, Implement is identical to DNA barcoding and is available via a pop-up window.

<u>Stage 3 – Analyze</u>: MAP guides the user through the four analytical steps required to generate a list of the species detected in each sample: 1) Import sequence data; 2) Select reference library; 3) Perform bioinformatic analyses; and 4) Evaluate technical/biological summaries. Its first two steps mirror those for barcoding, but data analysis is driven by MAP. Its capacity to deliver taxonomic information on the OTUs recovered from a sample depends heavily upon the completeness of the DNA barcode reference library for the study area. OTUs that find a strong match (> 97.7% similarity) to BINs/species in the library can be assigned to this taxon with high confidence, but those showing deeper divergence may reflect newly encountered species, sequencing artefacts (e.g., chimeras), or NUMTs. As with the DNA Barcoding workflow, bioinformatics parameters can be tuned by the user.

***Reference Libraries***: Both the DNA barcode and metabarcode workflows require access to a reference sequence library to assign taxonomy and BIN identifiers to OTUs. In the case of DNA barcoding, this library also facilitates the delineation of target from non-target sequences. In the case of metabarcoding, the reference library is required to assign the OTUs recovered from a bulk sample to their source species. BOLDistilled, ONTOLOGY’s default COI library, is updated quarterly and currently includes reference COI sequences from over 1.1M species of animals, bacteria, fungi, plants, and protists [25]. However, ONTOLOGY can also accept user-specified libraries that include a geographic or taxonomic subset of these records. The user assigns a name to the library that is imported by ONTOLOGY as a SINTAX-formatted *.fasta file [26].

***Custom Runs***: ONTOLOGY supports custom experimental designs—that is, those not using ONTOLOGY kits—for both of its core workflows. For custom Runs, users must enter primer and UMI sequences into the metadata and import relevant parameters and reference libraries. Aside from the supply of this additional metadata, Custom Runs are identical to kit-based Runs.

### 3. Technical Demonstration

**Sample and data collection**: Five Malaise traps were deployed in Muogamarra Nature Reserve, New South Wales, Australia, from December 3–29, 2024. Specimens captured by these traps were harvested on December 21 and December 29, producing a total of ten bulk samples. Details regarding sampling and DNA extraction were previously reported by Prosser *et al.* [10]. These samples were used to produce data required to demonstrate the effectiveness of ONTOLOGY.

**DNA Barcoding:** Ten ONTOLOGY PCR plates (AR-01) were used to process 950 DNA extracts following the standard ONTOLOGY protocol. The ten initial plates were selected based on high sequencing success among wells. The prepared library was sequenced on a Flongle using an ONT MinION Mk1D sequencer. Although these specimens have been previously analyzed [10], the data from this specific sequencing run are original to this manuscript.

**Metabarcoding:** One ONTOLOGY symmetrical metabarcode plate (AR-01-SYM) was used to process the ten samples, each of which included eight technical replicates. Half of the remaining 16 wells were used as positive control wells containing a synthetic control, while the other eight were negative control wells that only contained PCR cocktail. This sequencing run was previously analyzed [24].

**Reference Library**: The July 2026 BOLDistilled library [25] was employed in the analysis. Among the global records, it included 3,013 BINs derived from the barcode analysis of 22,856 specimens collected at Muogamarra between 2015 and 2025.

## RESULTS

**Barcode Results**: Analysis of the ten ONTOLOGY AR-01 plates on a Flongle generated 507,305 reads. One or more OTUs were recovered from 931 of the 950 DNA extracts. Correspondence between the specimen’s ordinal assignment based on its morphology and the taxonomy assigned via BOLDistilled led to the recovery of a DNA barcode (i.e., ‘target’ sequence) from 887 specimens. In addition, non-target OTUs were recovered from 378 specimens. A total of 300,256 reads contributed to target barcodes while 23,891 reads contributed to non-target sequences (NTS), meaning that 63.9% of raw reads were retained. Of the 887 specimens that obtained a target barcode, 95% would have been obtained with only 8.7K assigned reads (about 9 per specimen; Figure S2A).

**Metabarcode Results**: Analysis of the metabarcode plate on a PromethION generated 179,806,945 reads, of which 147,291,152 were assigned to an OTU (81.9%). Individual samples yielded from 193 to 599 BINs, and the mean count was 373. Detailed analysis of this dataset in comparison with the DNA barcode dataset will be considered in a future study. Metabarcode analysis required 26M assigned reads (approx. 2.6M per sample) to detect 95% of the BINs present in the Run (Figure S2B).

## DISCUSSION

ONTOLOGY was designed after joint consideration of user needs and the capabilities of nanopore sequencing platforms. When design was initiated, ONT was the sole vendor of these sequencers, and it produced three flow cells (Flongle, MinION, PromethION) with a 150-fold difference in their mean COI sequence output and a 13-fold cost differential ($85–$1,140 USD). These flow cells defined the possible, while the user community provided guidance on the deployment of this sequencing capacity. Since the initiation of ONTOLOGY’s development, several firms have entered the nanopore market [27, 28] and at least one new platform (Qitan Tech) has capabilities similar to ONT [7].

<u>Barcode Analysis</u>: The Flongle Run analyzed in this paper returned a mean sequence depth of 348 reads per specimen, more than sufficient to obtain accurate consensus DNA barcode sequences from specimens in a highly multiplexed pool [12]. ONTOLOGY’s Barcode workflow was designed to employ the Flongle flow cell, but ONT will discontinue it in February 2027. As a result, most users will need to employ an alternative flow cell for barcoding, such as an ONT MinION or Qitan QCell-384, both of which can be washed for reuse [28, 29]. At London Calling 2026, ONT revealed plans for a low-throughput flow cell to replace the Flongle, but the QCell-384 from Qitan Tech is currently the nanopore flow cell best suited for DNA barcoding, though it is not widely available outside of Asia.

<u>Metabarcode Analysis</u>: While ONT flow cells easily deliver enough reads for DNA barcoding, metabarcoding requires deep sequencing to maximize species detection because species vary in abundance and body size. As well, differential amplification linked to varying primer-template mismatches enhances variation in amplicon counts. As a result, metabarcode studies require 1M reads per sample to maximize species recovery from bulk samples [29, 30]. Given this target, the PromethION is the sole ONT flow cell suitable for organismal metabarcoding, and it can only analyze one plate of DNA extracts at this read depth. Until now, metabarcoding studies have relied on Illumina sequencers because they deliver large numbers of reads. For example, a NovaSeq X run can generate 50 billion reads, but this platform comes with a constraint: it can only characterize short amplicons (< 200 bp). While Illumina’s MiSeq i100 can sequence full-length barcode amplicons, the cost ($100 per million reads) is 10-fold higher than with ONT. The acquisition of reads spanning the 658 bp barcode region of COI is desirable because it reduces exposure to NUMTs [31] and helps to ensure correct assignments for closely related species with low COI divergence. In the future, ONTOLOGY will incorporate MAP’s functionality to process Illumina sequences.

This study demonstrated ONTOLOGY’s capacity to support barcode and metabarcode workflows by analyzing ten bulk samples and 950 single specimens. When viewed from the perspective of biological insights per million reads, the differential efficiency of these two approaches is striking. The metabarcode analysis required 147M processed reads to reveal 1,501 BINs when queried against BOLDistilled. Many additional OTUs found no BIN match in the reference library. They could represent species missing from the library, NUMTs, or sequencing errors. If presumed to represent missing species, most can gain a taxonomic assignment at the ordinal level or better, but these ‘ghost species’ can only be validated by expansion of the reference library. Although metabarcoding delivered considerable information on species composition, it cannot yet provide abundance data. By comparison, barcoding delivers a more nuanced perspective on biodiversity. Based on 0.3M processed reads, barcode analysis revealed the presence and abundance of 292 unique BINs among the 887 target barcodes from the analysis of 950 specimens from one bulk sample. Moreover, secondary OTUs were recovered from over a third of the specimens, and their inspection revealed biologically significant associations. Because barcoding directly quantifies the abundance of each species, accumulation curves can be calculated to estimate taxon diversity, and abundance-based biodiversity metrics can be reported. By coupling the use of ONTOLOGY with nanopore sequencing, a specimen can be barcoded for less than $1. Because the specimen-centric perspective inherent in DNA barcoding offers many advantages, the use of metabarcoding should never be the lone protocol employed to quantify biodiversity.

By coupling nanopore sequencing [32] with ONTOLOGY, a highly functional barcode node can be established for roughly 20K USD, and a dual barcode/metabarcode node for roughly 50K USD (Table S1). Despite their modest cost, these nodes have the potential to generate considerable data. Presuming five runs per PCR machine for each of 250 days, a barcode node can process 237,500 specimens a year. Presuming triplicate analysis of each bulk sample, a node provisioned with a single P2 Solo could metabarcode 6,000 bulk samples in a year.

## CONCLUSIONS

ONTOLOGY currently supports barcoding and metabarcoding using the standard COI barcode region for the animal kingdom via its PCR kits, but additional primer sets will be introduced to support work on plants (ITS2 + rbcL) and fungi (ITS), among others. Future developments for the barcode workflow will also enable the joint analysis of several gene regions (e.g., COI + 18S) via multiplexed PCR to support phylogenetic placements and to aid the detection of species interactions. Primer sets will also be expanded to support the recovery of standard eDNA markers. By supporting both DNA barcoding and metabarcoding studies on diverse taxonomic groups, ONTOLOGY is well positioned to aid molecular studies of biodiversity in settings ranging from university and governmental laboratories to remote field stations.

## ABBREVIATIONS

BIN: Barcode Index Number; BIP: Barcode Inference Pipeline; BOLD: Barcode of Life Data System; COI: cytochrome c oxidase subunit I; eDNA: environmental DNA; GUI: graphical user interface; MAP: Metabarcoding Analysis Pipeline; NTS: non-target sequence; NUMT: nuclear mitochondrial DNA segment; ONT: Oxford Nanopore Technologies; OTU: operational taxonomic unit; PCR: polymerase chain reaction; UMI: unique molecular identifier.

## DECLARATIONS

### Ethics approval and consent to participate

Not applicable.

### Consent for publication

Not applicable.

## ACKNOWLEDGEMENTS

We thank Kevin Kerr, Spencer Monckton, Sameer Padhye, Nguyen Nguyen, Kristen McCabe, and Alexandra Dolezal for assistance with internal testing. We thank Emily Hartop, Ben Price, Jordan Beasley, Tomasz Mamos, Santiago Jaume, Juliana Morales, Marko Mutanen, Mikko Vallinmäki, Torbjørn Ekrem, Rodolphe Rougerie, and Avas Pakrashi for assistance with beta testing. Additional consultation with Dirk Steinke, Evgeny Zakharov, and Masha Kuzmina improved the software.

## Availability of data and materials

The datasets supporting the conclusions of this article, including the input data, pipeline outputs and the script used to produce Figure S2, are available in the figshare repository, https://doi.org/10.6084/m9.figshare.35094301. The raw ONT reads for the metabarcoding run are available in the NCBI Sequence Read Archive under BioProject accession PRJNA1496812.

## Availability and requirements

**Project name:** ONTOLOGY

**Project home page:** https://www.ontology.bio

**Archived version:** https://doi.org/10.6084/m9.figshare.35094301 (ONTOLOGY v1.01 for Windows, macOS and Linux)

**Operating system(s):** Windows 11, 64-bit (x64, Intel or AMD; ∼520 MB); macOS Tahoe 26 (Apple silicon, M1 or later; ∼640 MB); Ubuntu 24.04 LTS (x86_64; ∼430 MB). Windows 10, Intel-based Macs, Windows on ARM, Ubuntu 22.04 and earlier, and WSL are not supported.

## Programming language: Primarily Python

**Other requirements:** 10 GB RAM minimum (16 GB recommended); about 2.5 GB of disk space for the application, plus at least 8 GB more if the BOLDistilled reference library is used; administrator or sudo rights for installation. The application runs offline once installed, although an internet connection is needed to download BOLDistilled and send data to BOLD.

**License:** Proprietary licence, free of charge for research, educational and other non-commercial use; the source code is not provided. Full terms are given in the Supplementary Information (Software Licence and Privacy).

**Any restrictions to use by non-academics:** Commercial use requires prior permission from the Centre for Biodiversity Genomics, University of Guelph.

## Competing interests

The authors declare no competing interests.

## Funding

The development of ONTOLOGY was enabled by support to PDNH from the Gordon and Betty Moore Foundation (GBMF-12262), from the Government of Canada through Ontario Genomics and Genome Canada (OGI-208 and OGI-233), through a Major Science Infrastructure award (MSI 42450) from the Canada Foundation for Innovation, and through the Government of Canada’s New Frontiers in Research Fund (NFRFT-2020-00073). KAT was supported by a Mitacs Elevate Fellowship. The funders had no role in the design of the study, data collection and analysis, or preparation of the manuscript.

## AUTHOR CONTRIBUTIONS

PDNH: Conceptualization, manuscript draft, resource acquisition

EO: Software development, bioinformatics – design & implementation

SJ: Methodology – Freeze-drying protocols KAT: Project coordination, manuscript draft

RMF: Methodology – Molecular methods, data collection, Bioinformatics – COI sequence correction

SWJP: Bioinformatics – design

NWB: Bioinformatics – Contaminant removal, software testing SB: UI and graphic design

## AI USE

Claude Opus 5.5 was used to provide copyedit suggestions on the manuscript that were evaluated by authors.

## ADDITIONAL FILES

**Additional file 1 (.docx):** Supplementary Information. Production and longevity of the freeze-dried PCR plates (Figure S1), read depth required for barcode and BIN recovery (Figure S2), DNA barcoding and metabarcoding protocols, equipment costs (Table S1), and the ONTOLOGY software licence and privacy terms.

## SUPPLEMENTARY INFORMATION

### PCR PLATES

The balance of this section summarizes the key processes involved in producing the freeze-dried PCR plates which generate tagged amplicons for use in the DNA barcode/metabarcode workflows.

**PCR Cocktails**: Each well was filled with a 10.5 µL cocktail containing: 6.25 µL of 10% trehalose (Fluka Analytical), 1.62 µL of molecular grade water (Hyclone), 1.25 µL of 10X PlatinumTaq buffer (Thermo Scientific), 0.625 µL of 50 mM MgCl_2_ (Thermo Scientific), 0.0625 µL of 10 mM dNTPs (KAPA Biosystems), 0.625 µL of 2 μM forward primer (IDT), 0.0125 µL of 100 μM reverse primer (IDT), and 0.06 µL of 5 U/μL PlatinumTaq (Thermo Scientific).

**PCR Primers**: ONTOLOGY has one primer option for DNA barcoding at launch, which is designed for arthropods (AR-01). A kit with symmetrical primers (AR-01-SYM) is available to optimize sequence recovery in metabarcoding studies. The AR-01 primers target the 658 bp barcode region of the cytochrome *c* oxidase 1 gene (Hebert et al., 2003), and include a cocktail of insect primers LepF1/LepR1 and general primers LCO1490/HCO2198 [33, 34]. Though designed for arthropods, these primers are effective across a broad range of taxa including annelids, molluscs, nematodes, and even bacteria [35]. The AR-01 primers (5’–3’) are:

C_LepFolF = RKTCAACMAATCATAAAGATATTGG

C_LepFolR = TAAACTTCWGGRTGWCCAAAAAATCA

Visit www.ontology.bio for information on the availability of new primer sets.

**Unique Molecular Identifiers**: ONTOLOGY’s AR-01 DNA barcoding plates employ an asymmetrical tagging system which uses the 16 bp UMI sets developed by PacBio. Extensive validation studies have established that these UMIs do not interfere with PCR amplification across diverse taxa [34] and that they are effective with Oxford Nanopore [8]. The use of such long UMIs aids the assignment of each read to its source specimen while also being robust to a modest error rate. Symmetrical tagging does have the advantage that only one UMI needs to be recognized to assign a read to its source specimen, and as such it produces a 10% increase in assignment success in trials. As it is expensive to implement at large scale—it would require 1,920 different primers to discriminate the 960 wells in ten plates versus the 106 needed for asymmetric tagging—it was not adopted for DNA barcoding given that far more reads are generally obtained than are needed. However, the AR-01-SYM kit provides a symmetrical tagging option which is optimal for metabarcoding studies that involve the analysis of just 96 samples in a Run.

**Freeze Drying**: Once PCR cocktails were dispensed into 96-well plates, they were processed in a Freezone Stoppering tray dryer (Labconco **#**794801300) coupled with a 6 Litre Console freeze dryer (Labconco #700611000). Twenty plates containing liquid PCR cocktails were placed on the shelves of the tray dryer and the shelf temperature was lowered to −40 °C over the course of two hours without vacuum. Full vacuum (0 mbar) was then applied, and primary drying was initiated (90 min at −40 °C) before the shelf temperature was gradually increased to 0 °C over ∼1 hour. After the plates were at 0 °C for 15 minutes, secondary drying was initiated. This involved a gradual increase in the shelf temperature to 22 °C over ∼1 hour, followed by a hold at this temperature for a second hour. The vacuum was then slowly released, allowing the chamber to return to ambient pressure within 10 minutes. The freeze-dried plates were removed, immediately heat sealed to prevent the ingress of moisture, and placed in storage at 4 °C until distribution.

**Longevity of PCR Reactions**: Thirty identical PCR plates were freeze-dried in February 2025. Half were held at 22 °C (i.e., room temperature) and half at 4 °C. Amplification success was assessed after 1 week, 1 month, 2 months, 6 months, and 1 year. At these times, two plates from each thermal regime were haphazardly selected for analysis. To initiate PCR, 10.4 μL of molecular grade water was added to each well, and the four plates were then set aside for 10 minutes to allow dissolution of the PCR pellets. A 2 µL aliquot of a standard DNA extract (0.1 ng/μL from *Galleria mellonella*) was then added to each well before the plate was sealed and amplified using a standard thermocycling regime (see <u>DNA Barcoding</u> section below). Following PCR, amplification success was assessed by scoring bands on an E-Gel and by sequencing the amplicons on a Flongle. Plates held at 4 °C delivered nearly 100% amplicon recovery for the full year while those at room temperature showed similar success for the first two months, but recovery dropped to 5% after six months and to 0% after a year.

**Fig. S1.**
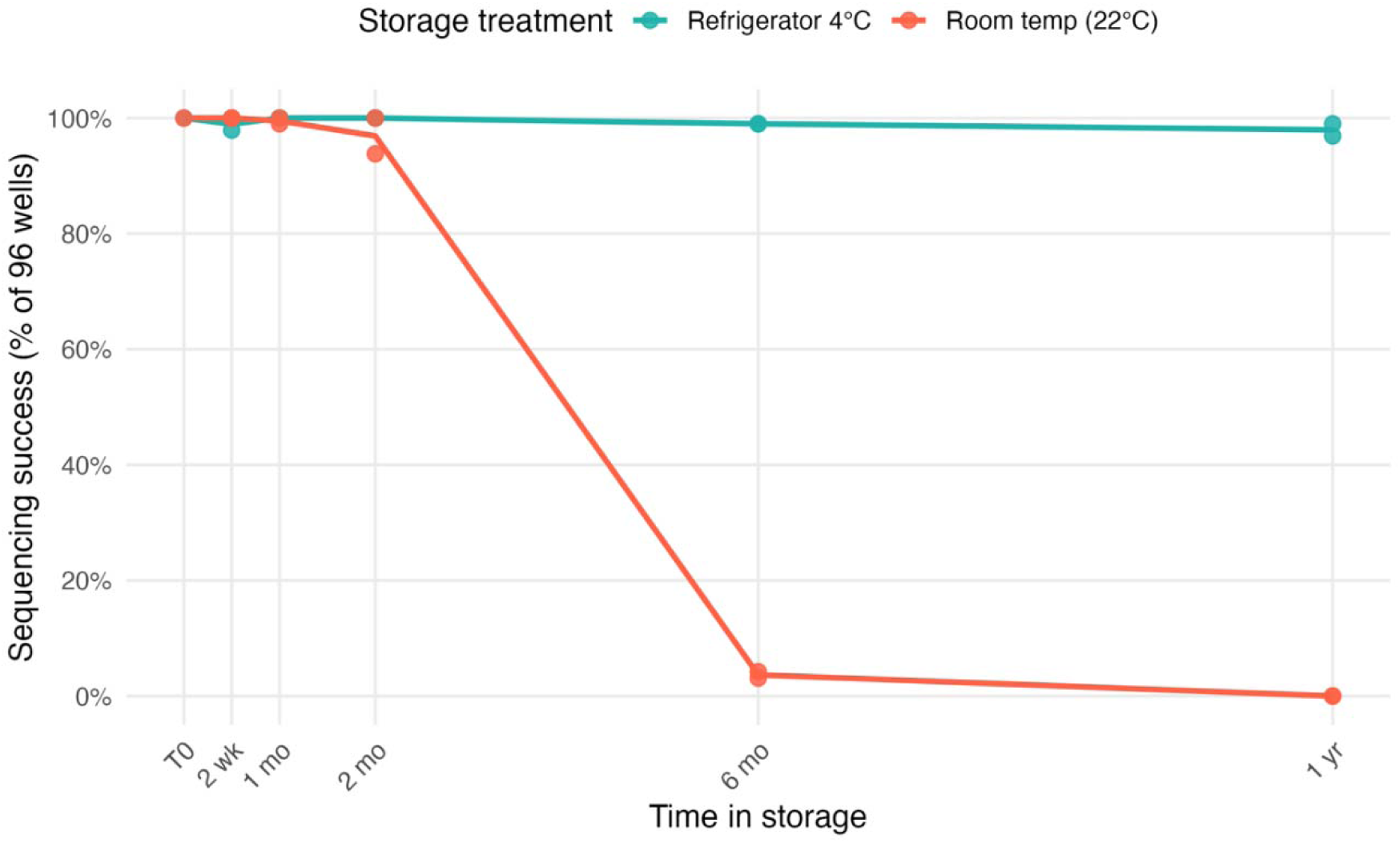
Longevity Test Results. The two temperature treatments are shown in separate colours.

**Fig. S2.**
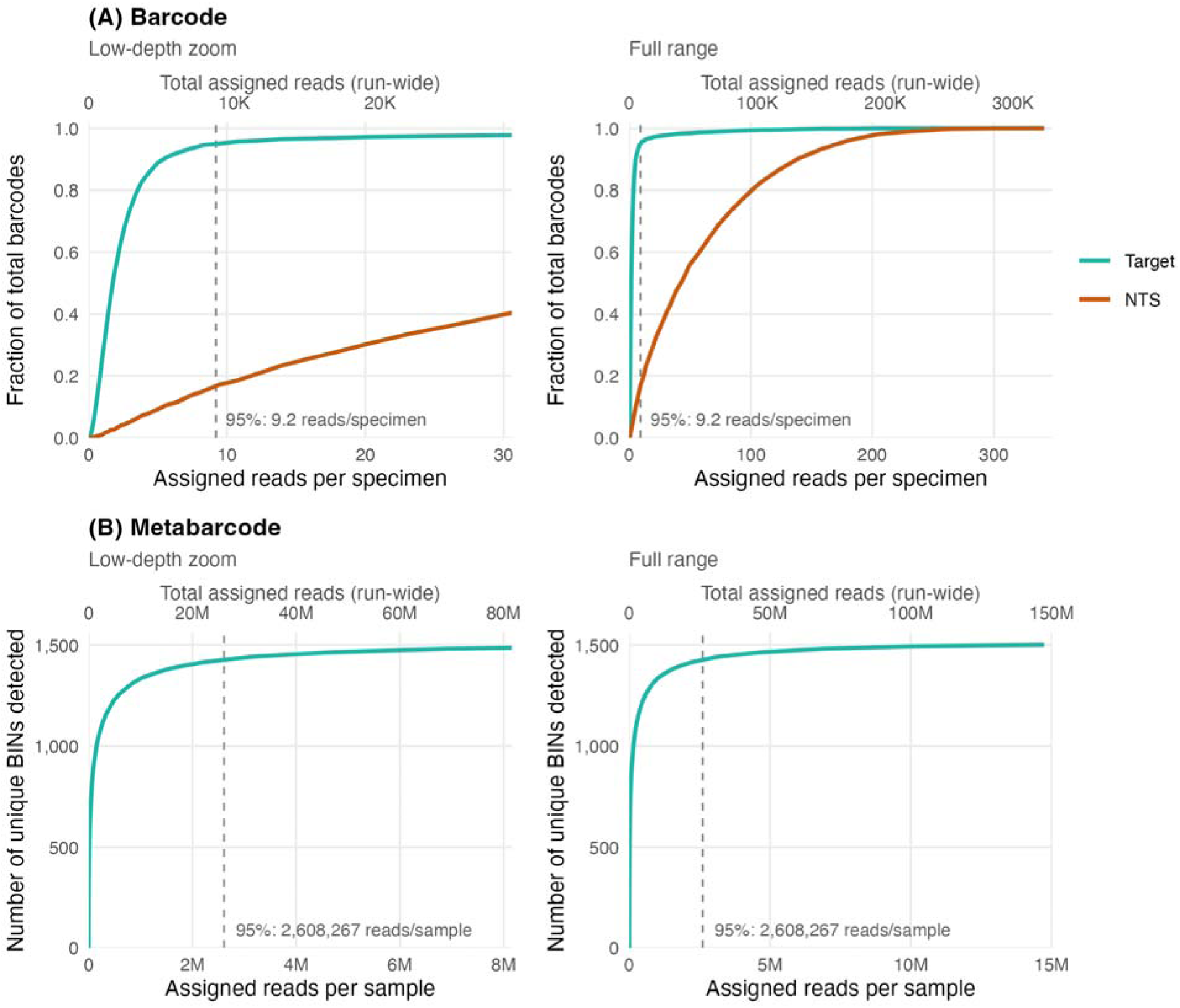
Analysis of read depth required to recover target barcodes and BINs in a Run. Vertical dashed lines indicate the point at which 95% recovery of (A) target barcodes and (B) metabarcode BINs is achieved. Non-target sequences—evidence of species interactions, NUMTs, or contamination—accumulate much more slowly than target barcodes as read depth rises. Curves show the mean of 20 random subsamples of assigned reads, drawn without replacement.

## DNA BARCODING

Barcode studies typically analyze a spectrum of taxa which differ in size, creating variation in concentration among their DNA extracts. The coupling of this variation with differing mismatches between templates and primers sets the stage for massive differences in amplicon concentrations after PCR and, hence, great variation in the read counts linked to each well following pooling. This variation was curbed by combining a high number of PCR cycles (i.e., 45) with limiting primer concentration. This protocol minimizes variation in amplicon concentrations among wells because primers are depleted, albeit after a varied number of cycles depending on template concentrations and primer-template binding efficiency. The effectiveness of this strategy has been demonstrated even when the pool of amplicons derives from 100,000 specimens [35]. Despite these steps to reduce well-to-well variation, some variation in amplicon concentration is inescapable and it is enhanced by the stochastic passage of amplicons through the nanopores. When success in barcode recovery was evaluated, it was found that it reached an asymptote when the number of assigned reads reached roughly 10× the number of specimens that donated amplicons to the pool. Flongle flow cells deliver a mean of 590K reads (SE = 31K), so they easily exceed the target read depth, even when specimens from ten plates are pooled. The following thermocycling regime is recommended for AR-01 plates:

94°C for 2 min; 5 cycles of [94°C for 40 sec, 45°C for 40 sec, 72°C for 1 min]; 40 cycles of [94°C for 40 sec, 51°C for 40 sec, 72°C for 1 min]; 72°C for 2 min.

**Barcode Inference Pipeline**: ONTOLOGY uses BIP as its bioinformatics backend for the DNA barcode workflow. BIP is designed to recover the target barcode sequence for each specimen that enters analysis, but it also retains any secondary OTUs and categorizes them as non-target sequences (NTS). Selection of the barcode sequence gains complexity when divergent sequences are recovered from a specimen. Some of this variation can arise from contamination, but other sequences derive from endosymbionts, parasites, parasitoids, or dietary components in the target specimen. BIP excludes chimeric [36] and low-quality sequences before it assigns each read to its source well by evaluating its UMIs. It subsequently trims primers and carries out two rounds of cluster analysis to delineate the OTUs represented among the sequences from each well. It then resolves sequencing and PCR error by constructing a consensus sequence for each OTU represented by two or more reads. This read requirement is based on extensive evaluation of the impact of sequencing error on the fidelity of sequences recovered by ONT [30]. BIP then scans the OTUs for NUMTs [31] and excludes them from further consideration. Finally, OTUs that contain indel errors in homopolymer tracts are corrected. Once this last phase of data validation is complete, BIP recognizes the dominant OTU as the barcode for the specimen when its taxonomic assignment determined through comparison with the reference library matches that based on morphology. Because required data volumes are modest (< 1M reads), ONTOLOGY’s implementation of BIP runs effectively on a laptop provisioned with 16 GB RAM and a modern multi-core CPU, typically completing analysis in less than two hours. Prosser et al. [17] provide a detailed discussion of BIP and its performance.

## DNA METABARCODING

DNA metabarcoding typically involves the analysis of a single plate with most of its wells dedicated to the amplification of DNA extracts from samples while the others are positive or negative controls. The high template diversity in DNA extracts from bulk samples favours the formation of chimeric sequences during PCR and their incidence rises with the number of cycles. To constrain their formation, the metabarcoding workflow employs 30 PCR cycles. The following thermocycling regime is recommended when AR-01 or AR-01-SYM plates are used for metabarcoding:

94°C for 2 min; 5 cycles of [94°C for 40 sec, 45°C for 40 sec, 72°C for 1 min]; 25 cycles of [94°C for 40 sec, 51°C for 40 sec, 72°C for 1 min]; 72°C for 2 min.

**Metabarcode Analysis Pipeline**: ONTOLOGY uses MAP as its bioinformatics backend for metabarcoding. As its components and functions are fully described by Prosser et al. [10], only a brief overview is provided here. The first seven steps in MAP mirror those in BIP, but the final step leads not to the selection of a single OTU but instead to a list of the BINs/species linked to the OTUs recovered from each sample. Because a PromethION Run can exceed 100M reads, analysis requires a desktop computer with ≥32 GB of RAM and sufficient storage. Users can also use ONTOLOGY’s plates with stand-alone MAP on a high-performance computing cluster. ONTOLOGY is currently only available as a downloadable application, but an online version will be developed to facilitate the analysis of metabarcoding Runs in the cloud.

## EQUIPMENT

ONTOLOGY makes it possible to develop a fully functional DNA barcode analytical facility for $20K USD and a facility with both barcode and metabarcode capability for roughly 2.5× as much. **Table S1** provides an overview of the equipment required to carry out the molecular components of the DNA barcoding and metabarcoding workflows.

**Table S1.**
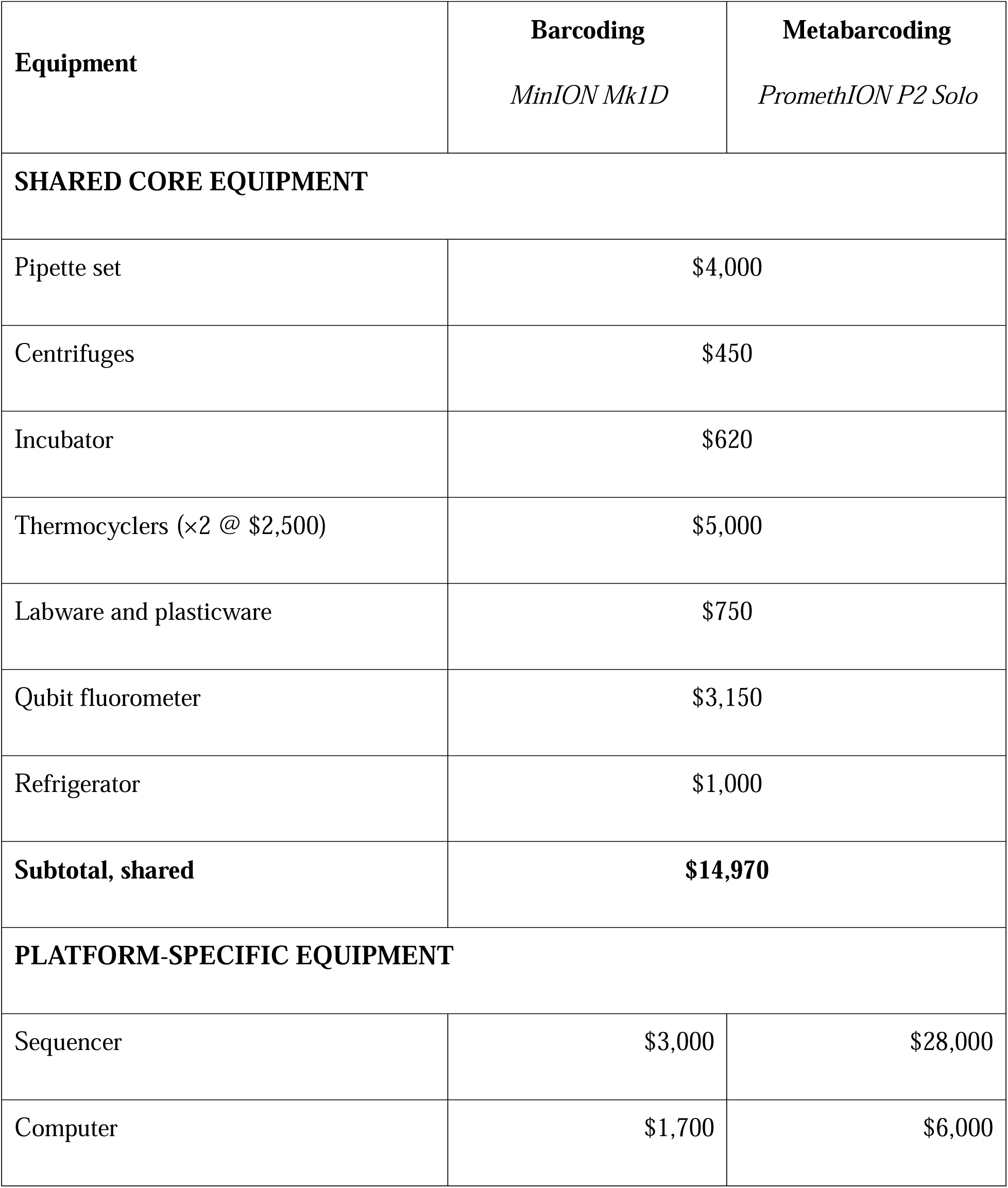
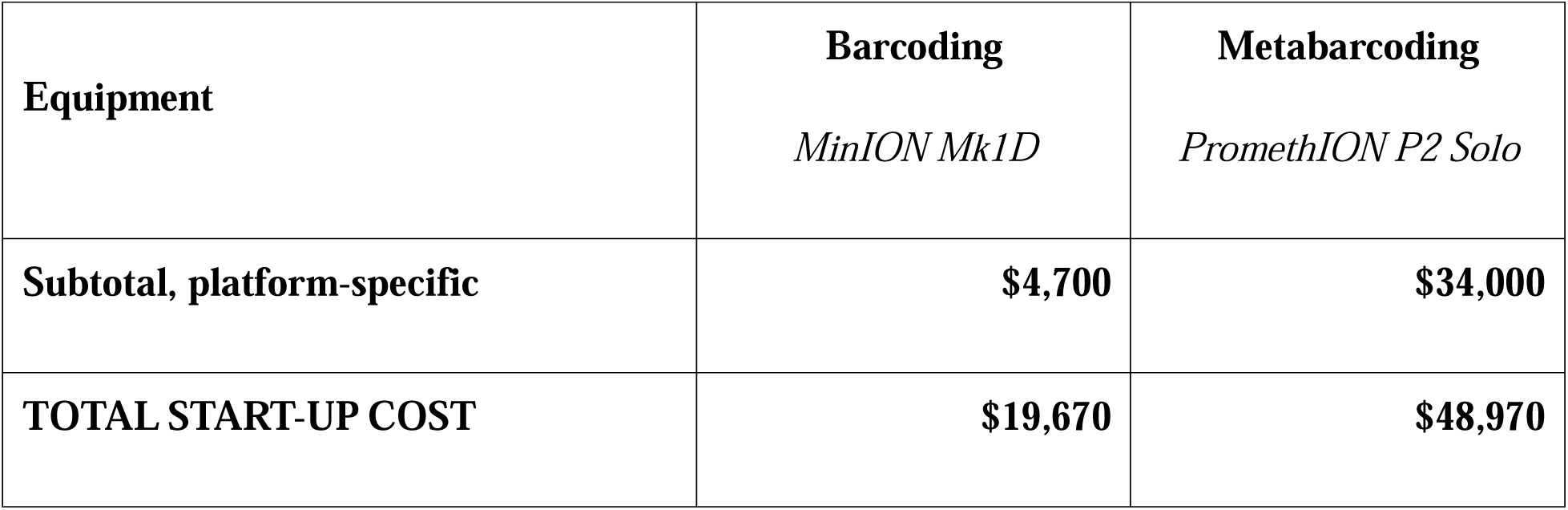
Equipment required to establish an ONTOLOGY barcoding and/or metabarcoding lab. Costs are approximate, in USD, and reflect prices in August 2026 in the Canadian market.

| Equipment | **Barcoding** | **Metabarcoding** |
|---|---|---|
|  | *MinION Mk1D* | *PromethION P2 Solo* |
| **SHARED CORE EQUIPMENT** |  |  |
| Pipette set | \$4,000 |  |
| Centrifuges | \$450 |  |
| Incubator | \$620 |  |
| Thermocyclers (×2 @ \$2,500) | \$5,000 |  |
| Labware and plasticware | \$750 |  |
| Qubit fluorometer | \$3,150 |  |
| Refrigerator | \$1,000 |  |
| **Subtotal, shared** | **\$14,970** |  |
| **PLATFORM-SPECIFIC EQUIPMENT** |  |  |
| Sequencer | \$3,000 | \$28,000 |
| Computer | \$1,700 | \$6,000 |
| Equipment | **Barcoding** | **Metabarcoding** |
|  | *MinION Mk1D* | *PromethION P2 Solo* |
| **Subtotal, platform-specific** | **\$4,700** | **\$34,000** |
| **TOTAL START-UP COST** | **\$19,670** | **\$48,970** |

## SOFTWARE LICENCE AND PRIVACY

ONTOLOGY is provided by the Centre for Biodiversity Genomics, University of Guelph (“CBG”), for research and educational use. By downloading or using the software you agree to the following terms.

**1. Licence grant.** CBG grants you a free-of-charge, non-exclusive, non-transferable licence to install and use ONTOLOGY on computers you control, for research, educational and other non-commercial purposes. For commercial use, please contact us first.
**2. Restrictions.** You may not sell, rent, sublicense or redistribute the software or its installers; modify, decompile, disassemble or otherwise reverse engineer it; use it to build a competing product; or remove any copyright or attribution notices. The source code is proprietary and is not provided.
**3. Ownership.** ONTOLOGY, including all updates, remains the property of CBG and the University of Guelph. This licence does not transfer any ownership rights to you.
**4. Third-party components.** The software includes open-source components that are distributed under their own licences, which continue to apply to those components.
**5. Citation.** If you use ONTOLOGY in work that you publish, we ask that you cite the software as indicated in its documentation or About dialog.
**6. Updates.** CBG may release updates at its discretion. Updates are covered by these same terms unless stated otherwise.
**7. No warranty.** The software is provided “as is”, without warranty of any kind, express or implied, including any warranty of merchantability, fitness for a particular purpose, accuracy of results or non-infringement. You use it at your own risk.
**8. Limitation of liability.** To the fullest extent permitted by law, CBG and the University of Guelph are not liable for any loss or damage, direct or indirect, arising from the use of or inability to use the software.
**9. Termination.** This licence ends automatically if you breach any of these terms. On termination you must delete all copies of the software.
**10. Governing law.** These terms are governed by the laws of the Province of Ontario and the federal laws of Canada applicable therein.

**Privacy.** The information you enter (name, email, institution, intended use, comments), your browser type and a one-way hash of your network address are stored in Canada and used only to understand who is using the software and to contact you about important updates. We do not sell or share this information. Contact to have your record removed.

## Notes

https://doi.org/10.6084/m9.figshare.35094301

### Competing Interest Statement

The authors have declared no competing interest.

## REFERENCES

1. Hebert PDN, Cywinska A, Ball SL, deWaard JR. Biological identifications through DNA barcodes. Proc Biol Sci. 2003;270:313–21. 10.1098/rspb.2002.2218.

2. Taberlet P, Coissac E, Pompanon F, Brochmann C, Willerslev E. Towards next-generation biodiversity assessment using DNA metabarcoding. Mol Ecol. 2012;21:2045–50. 10.1111/j.1365-294X.2012.05470.x.

3. Deiner K, Bik HM, Mächler E, Seymour M, Lacoursière-Roussel A, Altermatt F, et al. Environmental DNA metabarcoding: Transforming how we survey animal and plant communities. Mol Ecol. 2017;26:5872–95. 10.1111/mec.14350.

4. DeSalle R, Goldstein P. Review and Interpretation of Trends in DNA Barcoding. Front Ecol Evol. 2019;7:460426. 10.3389/fevo.2019.00302.

5. Leggett RM, Clark MD. A world of opportunities with nanopore sequencing. J Exp Bot. 2017;68:5419–29. 10.1093/jxb/erx289.

6. Cuber P, Chooneea D, Geeves C, Salatino S, Creedy TJ, Griffin C, et al. Comparing the accuracy and efficiency of third generation sequencing technologies, Oxford Nanopore Technologies, and Pacific Biosciences, for DNA barcode sequencing applications. Ecological Genetics and Genomics. 2023;28:100181. 10.1016/j.egg.2023.100181.

7. Thompson K, Prosser SWJ, Floyd R, Jafarpour S, Ozsahin E, Hebert PDN. Performance test of the QNome nanopore sequencer for arthropod DNA barcoding. bioRxiv. 2026;:2026.04.29.721586. 10.64898/2026.04.29.721586.

8. Hebert PDN, Floyd R, Jafarpour S, Prosser SWJ. Barcode 100K specimens: In a single nanopore run. Mol Ecol Resour. 2025;25:e14028. 10.1111/1755-0998.14028.

9. Srivathsan A, Loh RK, Ong EJ, Lee L, Ang Y, Kutty SN, et al. Network analysis with either Illumina or MinION reveals that detecting vertebrate species requires metabarcoding of iDNA from a diverse fly community. Mol Ecol. 2023;32:6418–35. 10.1111/mec.16767.

10. Prosser SWJ, Bard NW, Thompson KA, Floyd RM, Padhye S, Ozsahin E, et al. The Metabarcoding Analysis Pipeline (MAP): Simple, accurate, and flexible metabarcoding. bioRxiv. 2026. 10.64898/2026.07.22.740107.

11. Smith AM, Heisler LE, St Onge RP, Farias-Hesson E, Wallace IM, Bodeau J, et al. Highly-multiplexed barcode sequencing: an efficient method for parallel analysis of pooled samples. Nucleic Acids Res. 2010;38:e142. 10.1093/nar/gkq368.

12. Meyer D, Goettsch W, Spangenberg J, Stieber B, Krautwurst S, Hölzer M, et al. Unlocking the full potential of nanopore sequencing: tips, tricks, and advanced data analysis techniques. Nucleic Acids Res. 2026;54. 10.1093/nar/gkag023.

13. Sahlin K, Lim MCW, Prost S. NGSpeciesID: DNA barcode and amplicon consensus generation from long-read sequencing data. Ecol Evol. 2021;11:1392–8. 10.1002/ece3.7146.

14. Srivathsan A, Lee L, Katoh K, Hartop E, Kutty SN, Wong J, et al. ONTbarcoder and MinION barcodes aid biodiversity discovery and identification by everyone, for everyone. BMC Biol. 2021;19:217. 10.1186/s12915-021-01141-x.

15. Srivathsan A, Feng V, Suárez D, Emerson B, Meier R. ONTbarcoder 2.0: rapid species discovery and identification with real-time barcoding facilitated by Oxford Nanopore R10.4. Cladistics. 2024;40:192–203. 10.1111/cla.12566.

16. Hakimzadeh A, Abdala Asbun A, Albanese D, Bernard M, Buchner D, Callahan B, et al. A pile of pipelines: An overview of the bioinformatics software for metabarcoding data analyses. Mol Ecol Resour. 2024;24:e13847. 10.1111/1755-0998.13847.

17. Prosser SWJ, Thompson KA, Bard NW, Floyd RM, Ozsahin E, Hebert PDN. The Barcode Inference Pipeline (BIP): From Sequencer Output to DNA Barcodes. bioRxiv. 2026. 10.64898/2026.07.21.739876.

18. Jafarpour S, Floyd R, Prosser S. Production and Freeze-Drying of Indexed PCR Plates. protocols io. 11 2026. 10.17504/protocols.io.qpvrex7j2lmk/v1.

19. Klatser PR, Kuijper S, van Ingen CW, Kolk AH. Stabilized, freeze-dried PCR mix for detection of mycobacteria. J Clin Microbiol. 1998;36:1798–800. 10.1128/JCM.36.6.1798-1800.1998.

20. Yang S, Wen W. Lyophilized ready-to-use mix for the real-time polymerase chain reaction diagnosis. ACS Appl Bio Mater. 2021;4:4354–60. 10.1021/acsabm.1c00131.

21. Floyd R, Prosser S, Jafarpour S. DNA barcoding on Oxford Nanopore: multiplexing up to 24 x 96-well plates. protocols.io. 2023. 10.17504/protocols.io.rm7vzx6n4gx1/v1.

22. Mencius J, Chen W, Zheng Y, An T, Yu Y, Sun K, et al. Restoring flowcell type and basecaller configuration from FASTQ files of nanopore sequencing data. Nat Commun. 2025;16:4102. 10.1038/s41467-025-59378-x.

23. Ratnasingham S, Hebert PDN. bold: The Barcode of Life Data System (http://www.barcodinglife.org): BARCODING. Mol Ecol Notes. 2007;7:355–64. 10.1111/j.1471-8286.2007.01678.x.

24. Ratnasingham S, Wei C, Chan D, Agda J, Agda J, Ballesteros-Mejia L, et al. BOLD v4: A centralized bioinformatics platform for DNA-based biodiversity data. Methods Mol Biol. 2024;2744:403–41. 10.1007/978-1-0716-3581-0_26.

25. Prosser SWJ, Floyd RM, Thompson KA, Monckton SK, Hebert PDN. BOLDistilled: Automated construction of comprehensive but compact DNA barcode reference libraries. Mol Ecol Resour. 2025;25:e70043. 10.1111/1755-0998.70043.

26. Edgar R. SINTAX: a simple non-Bayesian taxonomy classifier for 16S and ITS sequences. bioRxiv 074161. 2016. 10.1101/074161.

27. Peng K, Yin Y, Li Y, Qin S, Liu Y, Yang X, et al. QitanTech nanopore long-read sequencing enables rapid resolution of complete genomes of multi-drug resistant pathogens. Front Microbiol. 2022;13:778659. 10.3389/fmicb.2022.778659.

28. Zhang T, Li H, Jiang M, Hou H, Gao Y, Li Y, et al. Nanopore sequencing: flourishing in its teenage years. Journal of Genetics and Genomics. 2024;51:1361–74.

29. Braukmann TWA, Ivanova NV, Prosser SWJ, Elbrecht V, Steinke D, Ratnasingham S, et al. Metabarcoding a diverse arthropod mock community. Mol Ecol Resour. 2019;19:711–27. 10.1111/1755-0998.13008.

30. Damm L, Khodami S, Rubel V, Molari M, Vink A, Stoeck T, et al. Community DNA outperforms eDNA metabarcoding for biodiversity assessments in the Clarion–Clipperton Fracture Zone. Metabarcoding Metagenom. 2026;10. 10.3897/mbmg.10.173793.

31. Hebert PDN, Bock DG, Prosser SWJ. Interrogating 1000 insect genomes for NUMTs: A risk assessment for estimates of species richness. PLoS One. 2023;18:e0286620. 10.1371/journal.pone.0286620.

32. González A, Fullaondo A, Odriozola A. Why are long-read sequencing methods revolutionizing microbiome analysis? Microorganisms. 2025;13:1861. 10.3390/microorganisms13081861.

33. Folmer O, Black M, Hoeh W, Lutz R, Vrijenhoek R. DNA primers for amplification of mitochondrial cytochrome c oxidase subunit I from diverse metazoan invertebrates. Mol Mar Biol Biotechnol. 1994;3:294–9.

34. Hebert PDN, Braukmann TWA, Prosser SWJ, Ratnasingham S, deWaard JR, Ivanova NV, et al. A Sequel to Sanger: amplicon sequencing that scales. BMC Genomics. 2018;19:219. 10.1186/s12864-018-4611-3.

35. Hoeksema BW, Roland, García-Hernández JE. A new host and range record for the gall Crab *Fungicola fageias* a symbiont of the mushroom Coral *Lobactis scutaria* in Hawai‘i. Pac Sci. 2018;72:251–61. 10.2984/72.2.7.

36. Edgar RC, Haas BJ, Clemente JC, Quince C, Knight R. UCHIME improves sensitivity and speed of chimera detection. Bioinformatics. 2011;27:2194–200. 10.1093/bioinformatics/btr381.

